# REVS: A New Open-Source Platform for High-Resolution Analysis of Rodent Wheel Running Behavior

**DOI:** 10.1101/2025.07.08.663718

**Authors:** James Bonanno, Ciara F. O’Brien, William B.J. Cafferty

## Abstract

**Background:** Rodent wheel running is widely used in neuroscience and preclinical research to assess locomotor function, recovery post-trauma or disease, circadian rhythms, and exercise physiology. However, most existing wheel-running systems offer limited metrics, lack flexibility in hardware, or require costly proprietary software, reducing their usefulness for detailed behavioral phenotyping—especially in models of injury or rehabilitation.

**New method:** We developed REVS (Revolution Evaluation and Visualization Software), a low-cost, open-source hardware and software platform for analyzing and visualizing rodent wheel running behavior. REVS captures wheel revolutions using Hall effect sensors and computes 13 day-level behavioral metrics along with detailed bout-level data. Users can interactively explore high-resolution temporal features and export data in Open Data Commons (ODC)-compatible formats. REVS supports customizable wheel types, facilitating use in animals with motor and/or sensory impairments.

**Results:** We validated REVS using a mouse model of partial spinal cord injury, where fine motor control is compromised. REVS detected impairments in 10 of 13 behavioral metrics post-injury, with varied recovery trajectories across measures. Principal component analysis revealed that recovery was closely linked to bout quality and intensity, rather than timing.

**Comparison with existing methods:** Unlike commercial and open-source systems, REVS offers more detailed metrics, customizable wheel compatibility, seamless blending with common vivarium hardware, integrated data visualizations, and ODC-compatible data export. It also supports flexible analysis across individuals and groups.

**Conclusions:** REVS provides a powerful, scalable tool for granular behavioral phenotyping in rodent studies, enhancing reproducibility and revealing insights into subtle locomotor changes associated with injury, recovery, and intervention.

**Highlights:**

- REVS enables detailed analysis of voluntary wheel running in rodents
- The platform combines low-cost hardware with open-source analysis software
- REVS computes 13 behavioral metrics across daily and bout-level timescales
- We identified a unique lesion and recovery profile after partial spinal cord injury
- REVS supports ODC-compatible data export for transparency and reuse

## 1. Introduction

Understanding rodent locomotion is foundational to a wide range of basic and preclinical research (Guo et al., 2019), from the study of motor systems (Grillner & El Manira, 2020; Lee et al., 2022) to circadian biology (Tam et al., 2015), cognition (Cotman & Engesser-Cesar, 2002), and aging (Kamel, 2003). Accurate measurement of locomotor behavior is especially critical for preclinical research into injury, recovery, and rehabilitation of the central nervous system, as it provides a window into the integrity and plasticity of motor circuits (Loy & Bareyre, 2019; Ward & Carmichael, 2020). As rodent models continue to evolve as translational tools for human disease, there is a growing need to quantify not only whether animals move, but how they move, with granularity that captures subtle shifts in behavior that might otherwise be missed.

Accordingly, in recent years neuroscience research has moved toward a more refined understanding of behavior, emphasizing the importance of tracking nuanced behavioral changes that can serve as early indicators of injury and hallmarks of disease progression and/or recovery. This growing emphasis has been driven by advancements including machine learning-based pose estimation tools like DeepLabCut (Mathis et al., 2018) and unsupervised behavioral clustering platforms like MoSeq (Lin et al., 2024; Weinreb et al., 2024), which have enabled investigators to extract detailed behavioral features from freely moving animals. These tools have highlighted the power of finely resolved behavior analysis to reveal distinct phenotypes not easily detectable through coarse, aggregate behavioral metrics (Akiti et al., 2022; Ashiquzzaman et al., 2025).

Despite these advances, there remains a gap in accessible, high-resolution systems for specifically analyzing voluntary locomotion, especially in the context of chronic in-cage wheel running, a widely used paradigm in rodent studies. Wheel running is a naturalistic, low-stress behavior that is frequently used to study voluntary exercise, circadian rhythms, and neurorehabilitation. However, most commercially available systems either limit data collection to simple metrics like distance or speed, or present cost and flexibility barriers that restrict their utility across research settings. Open-source and in-house solutions often address these limitations in part, but they typically lack detailed analysis tools or user-friendly visualization and export features. Additionally, previous open-source options rely on spinning-disk wheels, limiting the flexibility of experimental designs (Bivona & Poynter, 2021; Deitzler et al., 2022; Edwards et al., 2021; Godfrey et al., 2022; Mayr et al., 2020; Terstege & Epp, 2024; Zhu et al., 2021).

To address these limitations, we developed REVS: Revolution Evaluation and Visualization Software, a low-cost, open-source hardware and software platform for upright wheel running metric analysis and visualization. A central feature of REVS is its detailed quantification of locomotor behavior across multiple timescales. At the day level, REVS computes 13 distinct behavioral metrics, such as distance ran, as well as more nuanced measures like active cycle running proportion. These metrics are calculated automatically and can be visualized at both the group and individual animal level, stratified by custom groupings such as treatment condition – a significant advancement over available systems.

In addition to day-level analyses, REVS also performs high-resolution bout-level analysis. Each bout of running is defined by its temporal structure and annotated with metrics such as duration, speed, and distance. Users can explore these data via interactive millisecond-resolution bout timing plots or histogram-based visualizations of bout characteristics. Full day-level and bout-level data can be exported for statistical analysis or integration with other datasets. Critically, REVS is designed to promote open and transparent data reporting and sharing. All outputs can be exported in formats compatible with the Open Data Commons (ODC) framework (Torres-Espín et al., 2022), including optional metadata tables and automatically generated data dictionaries. This streamlines data sharing and enhances reproducibility.

In the present study, we applied REVS to a mouse model of partial spinal cord injury to determine whether this system could detect subtle behavioral impairments and recovery over time. By leveraging the full suite of REVS capabilities, we were able to identify distinct profiles of deficit and recovery across the 13 metrics and visualize bout-level changes that would be difficult to capture with traditional approaches. These results demonstrate that REVS is not only a tool for administering voluntary exercise, but also a powerful platform for behavioral phenotyping in injury and rehabilitation models.

## 2. Methods

### 2.1 Electronics and sensor building

Our movement detection system uses an Arduino Uno R3 microcontroller (Arduino A000066) and custom-prepared Hall effect sensors (Digi-Key 620-1330-ND) to detect the frequency and time of wheel revolutions.

#### 2.1.1 Preparing the Hall effect sensor

The Hall effect sensor is the first critical hardware component to be prepared (Fig. 1). The sensor (Fig. 1A) must be attached to a wire that transmits movement related signals to an Arduino control unit. The Hall effect sensor cables should be cut to a preferred length for your recording set up. Briefly, the excess plastic around the wire must be stripped to reveal the wiring (Fig. 1B). Next, the wire and the sensor must be tinned and then soldered together to complete the connection (Figs. 1C, D). The orientation of the Hall effect sensor is critical. Ensure that the sensor is top side up, designated by an imprinted capital A. Heat shrink should be applied over the soldered connections to protect them (Fig. 1E). For long-term durability, encase each Hall effect sensor in epoxy putty (Oatey 31270) (Fig. 1F).

**Figure 1.**
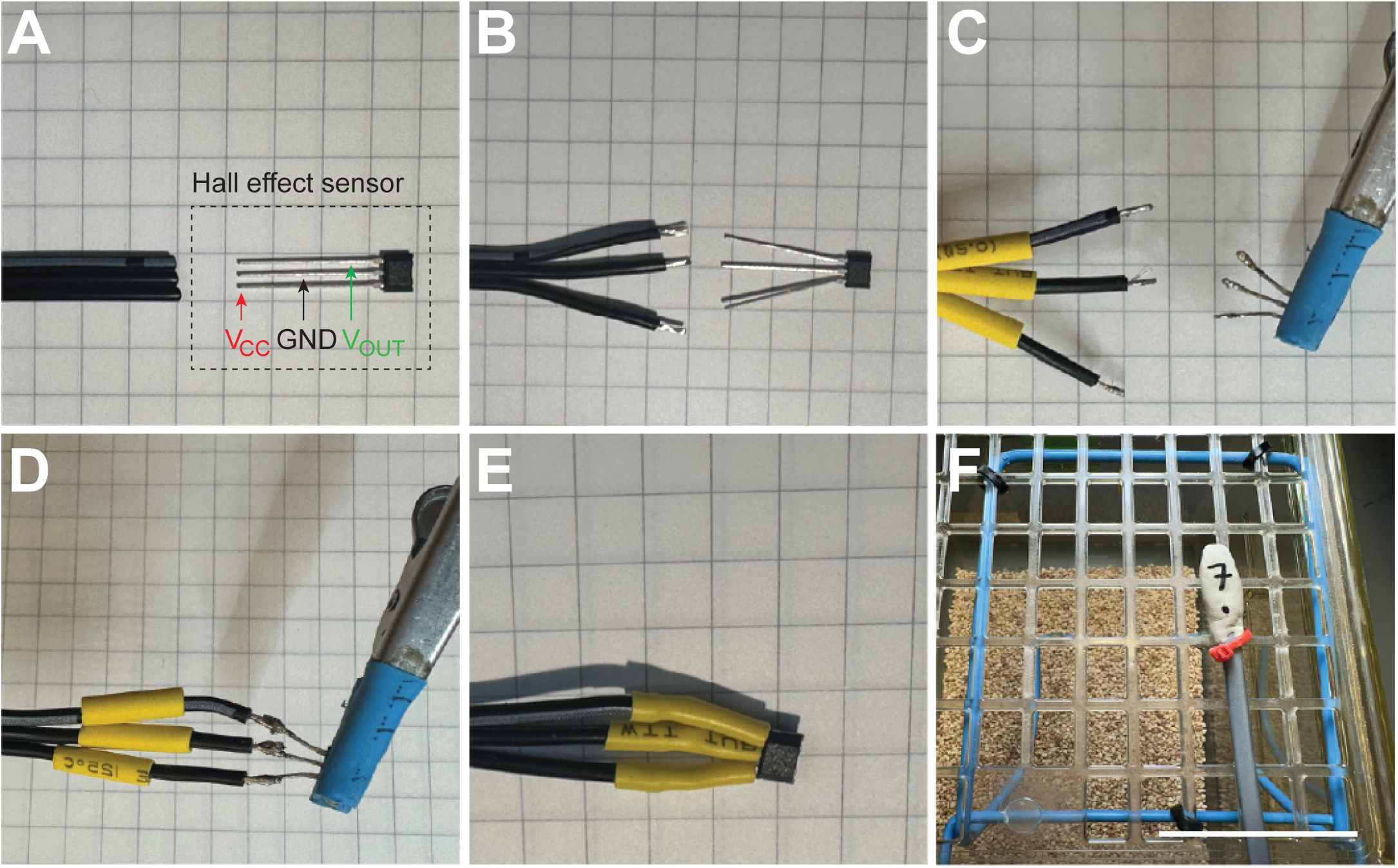
Hall effect sensor preparation. To prepare the Hall effect sensor, first align with 3-wire cable (**A**). Strip the end of the 3-wire cable ∼1 cm and separate the three prongs of the Hall effect sensor (**B**). Place heat shrink over each separated wire and tin each wire and each prong of the Hall effect sensor (**C**). Solder each of the three wires to the corresponding prong of the sensor (**D**). Move heat shrink over the soldered connections and apply heat to bind (**E**). To protect the sensor, encase in epoxy putty and secure to cage top in close proximity to running wheel (**F**). Graph paper (**A-E)** = 5 mm squares. Scale bar = 5 cm (**F**).

#### 2.1.2 Arduino electronics setup

We designed our Arduino to be suitable for voluntary movement detection of 8 singly housed mice. After creating the number of sensor cables suitable for your experiment, they must be connected to the Arduino control unit. Each sensor cable has 3 wires to be connected: VCC, a 5-volt input from Arduino; GND, ground input from Arduino; and V_OUT_, signal output from sensor to Arduino (Fig. 2A, 2B). Note which input pin each sensor cable is attached to and label the epoxy putty shield appropriately (Fig. 1F).

**Figure 2.**
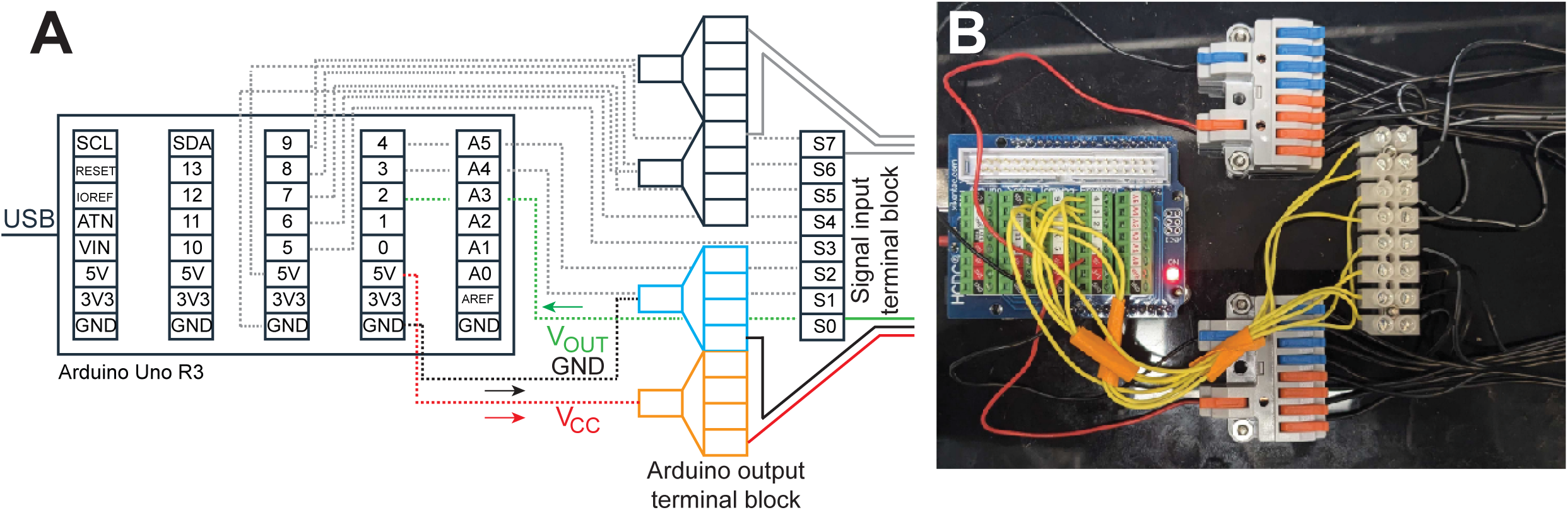
Overview of REVs hardware components. Arduino Uno R3 connectivity is schematized in panel **A** and demonstrated in panel **B**. Sensor cables are split and organized into one of two terminal blocks: the signal (V_OUT_) input terminal block, conveying movement signals to the Arduino, or the Arduino output terminal block supplying power (V_CC_) and grounding (GND) to Hall effect sensor. V_OUT_ wires (solid green line **2A**) from the sensor cable enter the signal input terminal block. Wires from the input signal terminal block (dotted grey lines **2A**, yellow wires **2B**) are then secured to individual pins on the Arduino. The provided Arduino software, by default, indexes 8 sensors starting at digital pin 2. Thus, for 8 sensors, they should be connected at digital pins 2, 3, 4, 5, 6, 7, 8, and 9. Single power (dotted red line **2A**) and ground (dotted black line **2A**) wires from the Arduino enter Arduino output terminal blocks (orange, power; blue, ground). Power and ground output is then split to supply individual sensor cables (black wires **2B)**.

### 2.2 Wheel construction and cage setup

Once sensor cables and the Arduino controller unit are prepared, in-cage running wheels need to be modified to track running behavior (Fig. 3).

**Figure 3.**
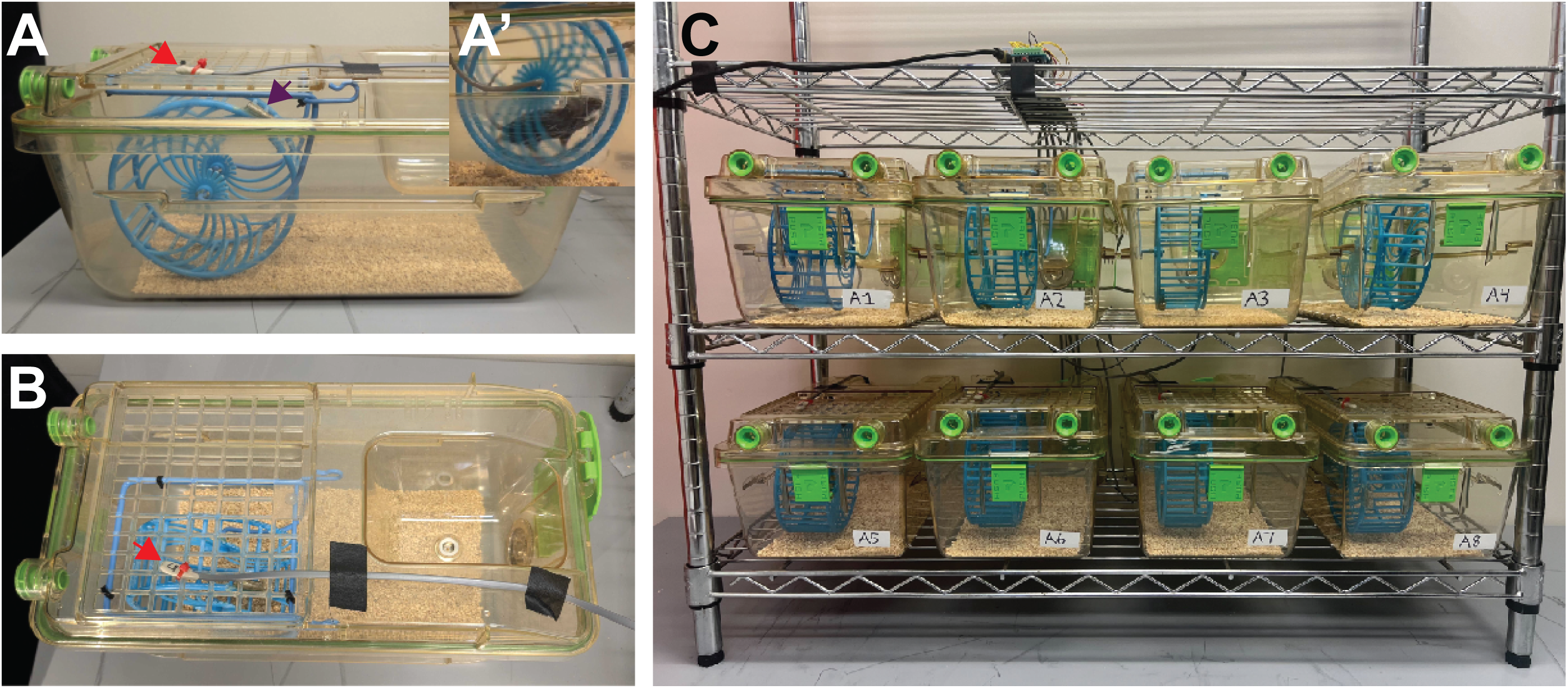
Overview of REVs wheel and cage setup. Wheels are shown suspended from cage top (**3A**, **3B**). Hall effect sensors (red arrowheads) connected to the Arduino interface directly with cage tops and detect magnets (purple arrowhead) as mice run (**A’**). An example of a fully equipped wheel-recording setup for 8 mice is pictured (**3C**).

#### 2.2.1 Wheel construction/modification

It is required that a wheel is selected that can fit within the bounds of the vivarium supplied cages. The exact dimensions and style of the wheel (i.e., runged versus flat bottom; sanded versus smooth; etc.) should be uniquely tailored to the experimental question. We used 12.5 cm diameter wheels (Walmart) and sanded the rungs on our wheels to provide more distinct sensory input. We left a subset of wheels with regular rung patterns for wheel-habituation and created a subset with a complex rung pattern for rehabilitation.

Additionally, the wheels must be modified to have two affixed magnets for Hall effect sensor activation (Fig. 3A, B). We used Gorilla 2-part epoxy (Amazon, 113351) to attach cylinder magnets (¼’’ x ¾’’, K&J Magnetics, #D4C-N52) to the outer circumference of the wheel. This process requires curing overnight after attaching the first magnet. Magnets should be placed 180 degrees apart and have their polarities in the same direction.

#### 2.2.2 Cage modification

Once the magnets are attached, the wheels must be placed in cages with two primary considerations: appropriate distance from the Hall sensor and unencumbered wheel rotation. We attached the axle from the Prevue Pet Products Wire-Mesh Wheel Toy (Prevue Pet Products SPV90011) to cage tops with zip ties to secure the wheel from above. The flexibility of the axle’s material facilitates the ability to adjust the wheel’s height within the cage. Alternatively, wheels can be upright from the cage-bottom. Note that the top of the wheel must be close enough to the top of the cage for the sensor to detect the magnets as the wheel rotates. Once the wheel placement in the cage is finalized, attach the sensor from 2.1 directly above the magnets (Fig. 3B) and secure with a zip tie. The magnet should be within approximately ∼1 cm of the sensor. To enable regular cleaning, ensure that the wheels are easily removable.

#### 2.2.3 Cage arrangement

Owing to the short range of the Hall effect sensor, cages can be arranged in close proximity around each Arduino setup (Fig. 3C). This cage arrangement is space efficient and allows for multiple setups in designated areas. Additionally, this arrangement keeps mice closely located to one another, which may blunt any negative effects of being singly housed.

### 2.3 Testing system and acquiring data

#### 2.3.1 Setting up the programming environment

To collect data on a computer or laptop, first download the Arduino IDE. For specific instructions, use the following Arduino support link: https://support.arduino.cc/hc/en-us/articles/360019833020-Download-and-install-Arduino-IDE. Each Arduino to be used must be connected to the computer via USB, and each Arduino can generally support up to 8 sensors simultaneously. Each Arduino must have the appropriate logging firmware uploaded through the Arduino IDE, which is provided in the GitHub repository, before it can be used. For each Arduino control unit, set up a folder with a unique name, such as “port_a,” and extract the given ZIP file into it. It should contain:

1. wheel-counter.jar, which is the executable JAR file that logs events from the Arduino and contains functions for log rotation and report generation.
2. start.bat, a batch file used to start wheel-counter.jar. It will need filling in once you extract it. It should contain the text “java -jar wheel-counter.jar log [ARDUINO COM PORT] [FOLDER NAME].log”. Fill in the parts in square brackets, where [ARDUINO COM PORT] is the port of the Arduino you would like to use, which can be determined by plugging in individual Arduino control units and checking which COM port corresponds to it in the Arduino IDE, and [FOLDER NAME] is the name chosen for the folder. The Arduino port can also be determined using Windows Control Panel.
3. rotate-log.bat, a batch file that will be called daily as a scheduled job by Windows to rotate the logs and generate a daily report. It also requires filling in. Its contents should be:

cd “[ABSOLUTE PATH TO THIS FOLDER]”
java -jar wheel-counter.jar analyze [FOLDER NAME].log
4. do-rclone.bat. Optionally, if using an rclone set up, this should contain “rclone copy ./ gdrive:wheel-counter/ --max-age 7d --include *.flog --include *.xlsx” depending on how the rclone remote is set up. If using an rclone set up, add to this folder a copy of the Windows 64-bit version of rclone.exe (available <u>here</u>). You will need to set up a token for Google Drive within your rclone installation. See here for instructions: https://rclone.org/drive/
5. pins2animals.txt, which should provide a new line-separated list of mappings from pin number to animal ID (of the form [PIN] -> [ANIMAL ID]), for use in daily report creation.

#### 2.3.2 Initializing data collection

Once all Arduino control unit directories are set up, the user will need to schedule a task to rotate and upload logs daily. We recommend using the Windows task scheduler for this process. To access on a Windows computer, hit the Windows Key + R and enter “taskschd.msc” into the run window that appears.

1. Select Task Scheduler Library in the left panel to bring up the general list of scheduled tasks.
2. Select Action > Create Task… in the menu bar.
3. Name your task, something like “Rotate Logs”
4. Select the Triggers tab in the dialog and hit “New…”

a. Begin the task on a schedule
b. Daily
c. Start tomorrow at the chosen time. This time should be during the expected inactive (day) cycle of the mice. For instance, if the mice are being run on a normal circadian cycle, 11:30 AM would be an appropriate time to rotate logs.
d. Recur every 1 days. Hit OK to commit the trigger.
5. For each Arduino directory set up select the Actions tab in the dialog and hit “New…”

a. Make sure “Start a program” is selected.
b. Click “Browse…” and navigate to the Arduino control unit folder directory in question.
c. Select “rotate-log.bat”
d. Hit OK to commit the action.
6. Hit OK to commit the task, then make sure to close the task scheduler, as the task may not run while it’s still open. To run the logging software, open a command line, navigate to the Arduino control unit folder directory you want to log for, and enter “start.bat”. It should begin logging immediately with a start message as the Arduino resets.

#### 2.3.3 Testing the wheel

Each sensor can be individually tested by briefly spinning the wheels once the logging window is open. Revolution events will appear in the console as they are logged with the appropriate pin number. We recommend labeling each sensor with the pin number that is reported in the logging software. If a sensor does not work, it is most likely an issue with a faulty connection between a sensor and the Arduino.

#### 2.3.4 Collecting and storing data

Whether uploaded locally on the device running wheel-counter, or to cloud storage, .flog files must be transferred to the computer that will run the REVS app in an appropriate folder structure. Groups of .flog files in which the pins correspond to the same set of animals must be together in their own folder.

Example: There are two Arduino setups, port_a and port_b, running simultaneously for 8 animals each, and each ran two distinct cohorts of animals at different times, cohort_1 and cohort_2. Each of these groups would need their own directory: Sample_experiment_data/port_a_1/*.flog, Sample_experiment_data /port_a_2/*.flog, Sample_experiment_data /port_b_1/*.flog, Sample_experiment_data /port_b_2/*.flog.

#### 2.3.5 Additional files

The final requirements to run REVS are input files to the GUI. These include a metadata file, a user created .csv file that contains grouping keys relevant to the experiment (i.e., animal ID, sex, treatment), and a JSON file that contains required export parameters. The export parameters are designed to be consistent with the Open Data Commons (ODC) framework to promote data sharing. We used a JSON file for the ODC for Spinal Cord Injury (ODC-SCI). A user-modified JSON file can be substituted to match the needs of particular experiments.

### 2.4 Graphical User Interface

In addition to providing a system for long-term wheel revolution data collection, we developed a user-friendly graphic user interface (GUI) which allows users to easily visualize their data and export it in formats compatible with ODC requirements. The GUI, titled Revolution Evaluation and Visualization Software (REVS) is composed of 4 major panels: Landing, Summary, Explore, and Export.

#### 2.4.1 Downloading and Launching REVS

REVS requires an installation of JAVA IDE later than version 8. To download and launch REVS, download the zip file provided at github.com/CaffertyLab/revs/. REVS can be launched by double-clicking the executable .jar file in the REVS directory.

#### 2.4.2 REVS panels and functionality

##### 2.4.3.1 Landing panel and settings dialog

Start by creating or loading a project on the landing page, where you will be required to input certain variables such as data input directories and wheel diameter (Fig. 4). These inputs include:

1. Dataset directory: Select a root folder contains data from each Arduino and cohort, stored separately into their own subfolders
2. Metadata file: Select a .csv file that contains information about your experiment such as animal IDs, their associated sensor pin path, and any grouping variables relevant for your experiment. An example metadata file is provided in an example project.
3. ODC requirements file: The JSON file, discussed above, which contains required export parameters.
4. Wheel circumference: A number input corresponding to the circumference of your wheels in meters.
5. Minimum Inter-Bout Duration: A number input corresponding to the minimum amount of time that must pass before the measurement of an initialized bout ceases.
6. Short Bout Threshold: A number input corresponding to the experimenter’s determination of what constitutes a “short bout”. This will directly impact the Short Bout Proportion measurement.
7. Minimum Revolution Threshold: A number input corresponding to the minimum number of wheel revolutions that must occur in a day to compute wheel metrics. We recommend a value of 100 to ensure reliable data.
8. Day Boundary: A time input in local time corresponding to the time at which data begins to count for the next day.
9. Start and End of Active Cycle: Time inputs corresponding to the timing of lights on/lights off for the given experiment.
10. Theme: A drop-down selection that allows users to choose between a light or dark GUI mode.
11. Palette: A drop-down selection that allows users to choose a color palette for the graphs in the explore panel.

**Figure 4.**
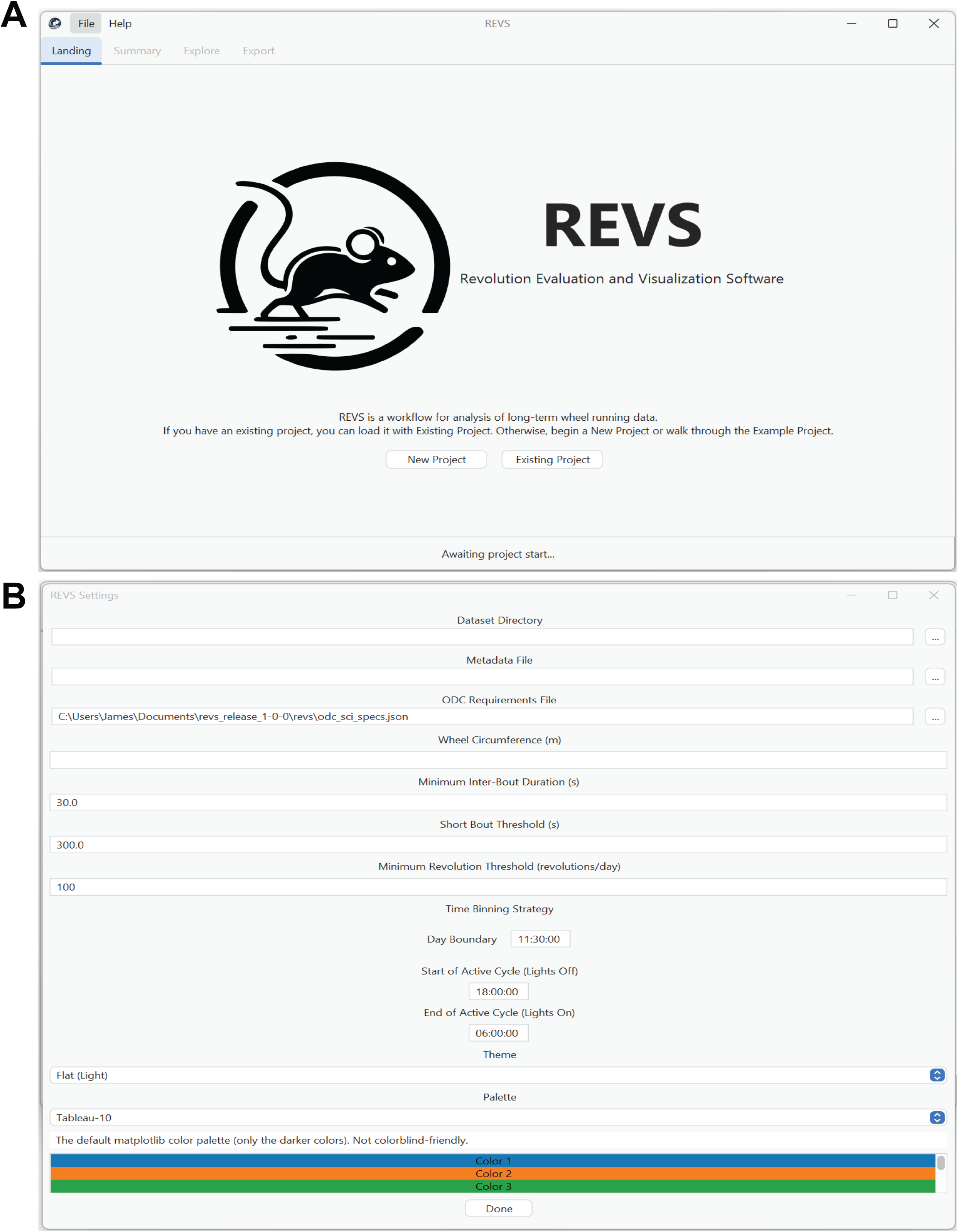
REVS landing page and settings. The landing page of the REVS GUI allows users to create a new project, load an existing project, or load an example project (**4A**). Upon selecting a new project, the user will be prompted to select input directories for data, metadata, and ODC settings. Additionally, the user will be prompted to input the wheel circumference for their experiment. The user can customize their experience by selecting a light or dark theme and a colorblind friendly palette (**4B**).

##### 2.4.3.2 Summary Panel

After creating your project, the summary panel will quickly summarize all animals in the entire dataset to verify group sizes (Fig. 5). The summary panel additionally provides a table of metadata which can be manually edited. If any necessary details are missing or incorrect (i.e., a missing sex or malformed date), the GUI will prompt users to correct them before moving on to data visualization. At any time, animals can be excluded from data visualization using the tree-diagram on the left. After the data is loaded and measurements are calculated, press “Begin Analysis”. You will then be able to use the Explore panel to visualize your data.

**Figure 5.**
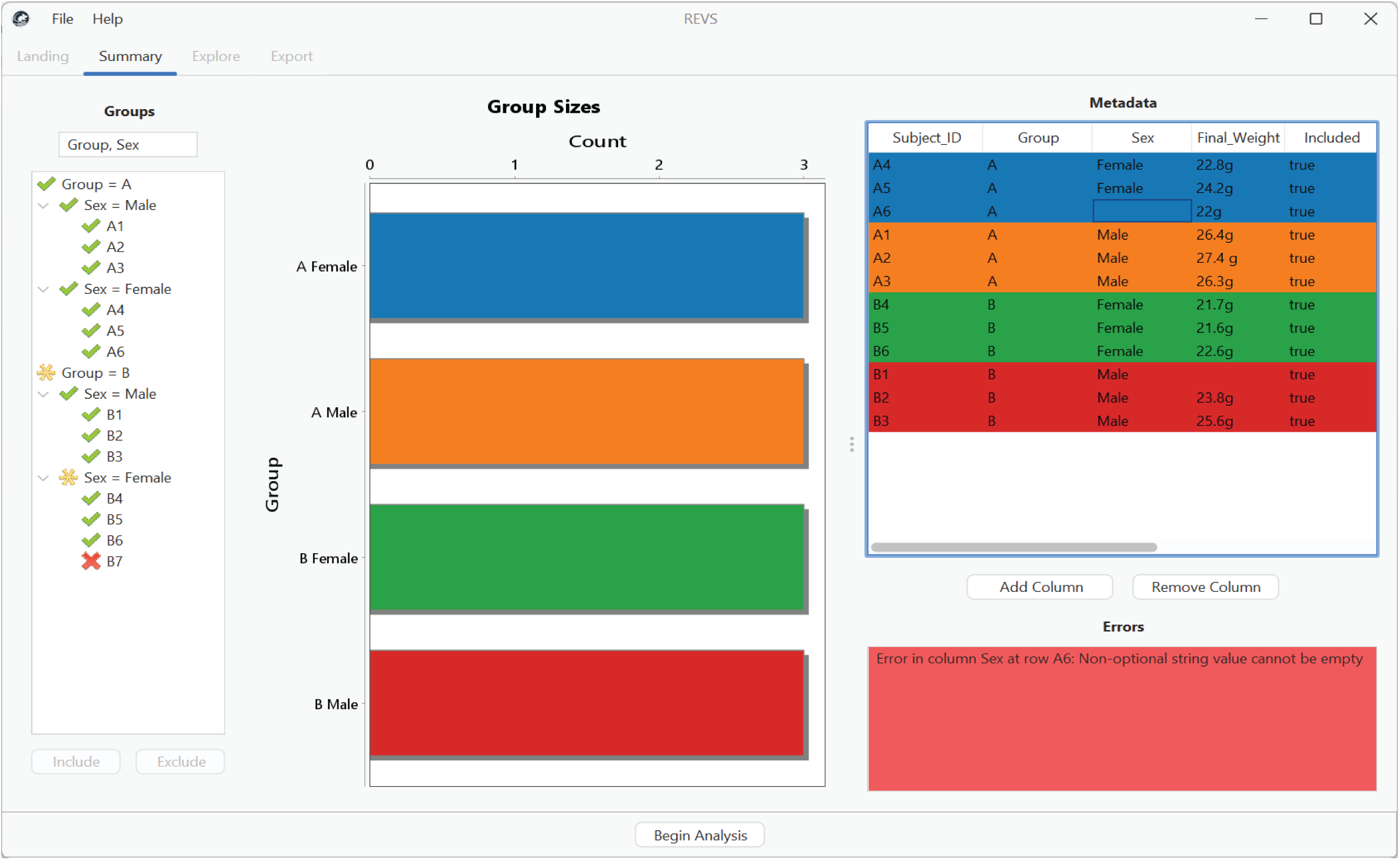
REVS summary panel. The REVS summary panel allows users to get a quick overview of their dataset. Drawing upon the user-created metadata file, the summary panel shows group sizes and allows the user to select grouping keys (middle). Animals can be excluded using the tree view (left). Additionally, the metadata file will be loaded in to be viewed as a table and can be modified (right). If there are any errors or missing values, the user will be prompted to fix them before running data analysis.

##### 2.4.3.3 Explore Panel

The explore panel (Fig. 6) allows users to visualize their data with four distinct graph types and thirteen distinct metrics. Additionally, the explore panel allows the users to select grouping variables and exclude individual animals, facilitating the ability to look at either whole or sub-groups of data. The Export Graph button saves a .csv file of all the data used to create the currently viewed graph, and the Export All button saves all data in a tidy format. The graph functionalities for the explore panel, and individual metrics, are detailed below.

**Figure 6.**
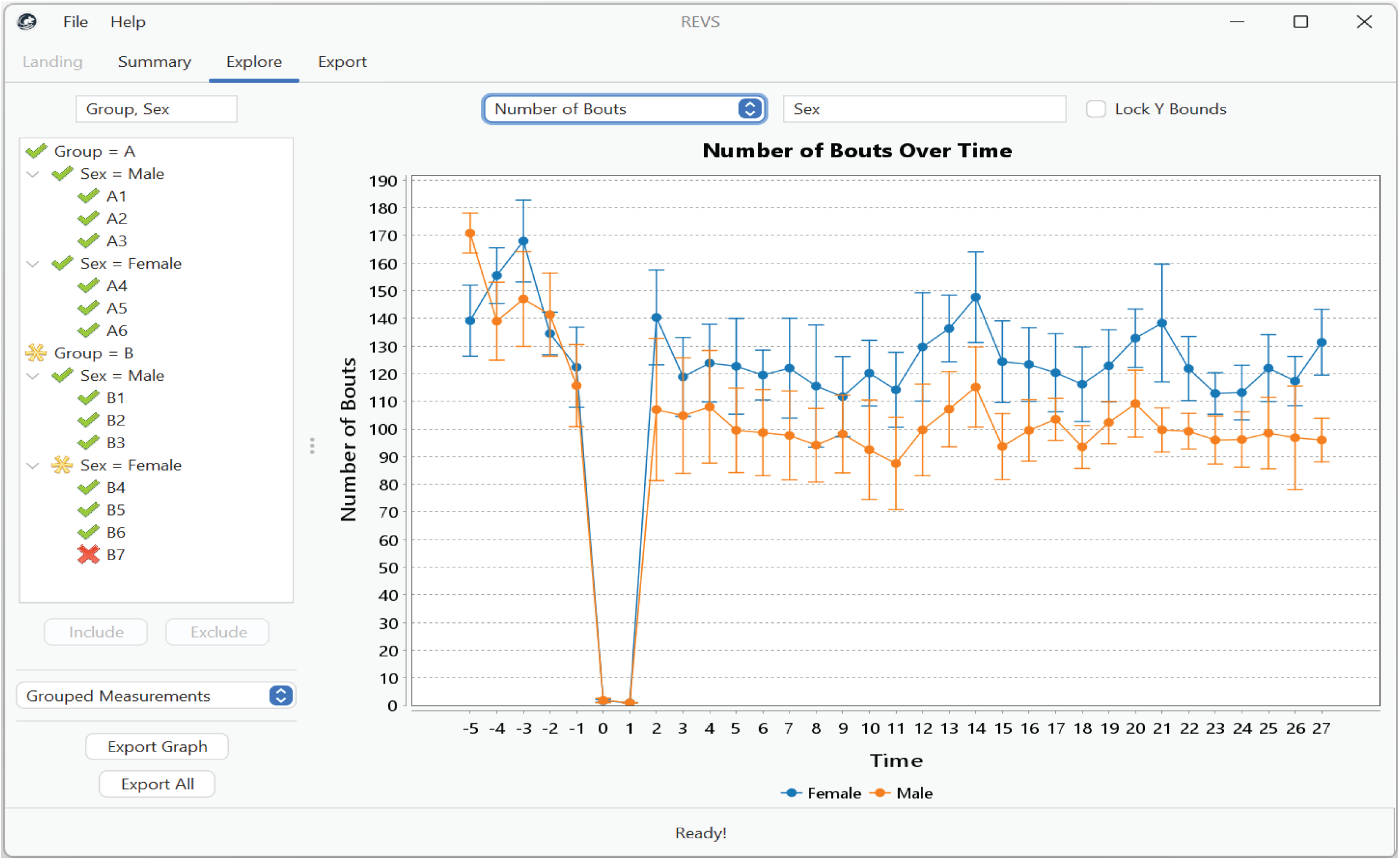
REVS explore panel. The REVS explore panel allows users to easily visualize wheel-running data from their experiment. Users are able to select from various graph types including grouped measurements and individual animal measurements. Users are also able to view day-level bout measurements for individual animals. The user can easily adjust grouping keys to look at measurements from different groups and can exclude animals using the tree view on the left. Critically, the user can adjust elements of the graphs such as axes by right clicking on the graph. Graphs and associated data, as well as all data, are exportable as CSV files using the Export Graph and Export All buttons respectively.

###### 2.4.3.3.1 Grouped Means and Individual Animal Data

The first two graphing functions allow users to visualize the group means and individual animal data for thirteen distinct metrics over the time course of the experiment. These thirteen metrics are comprised of 5 whole-day level metrics and 8 bout-level metrics (Figs. 7-8).

1. Revolutions – This metric measures the total number of recorded wheel revolutions over a single day.
2. Total Distance – This metric measures the total accumulated distance run over a single day.
3. Total Duration – This metric measures the total accumulated time running over a single day.
4. Overall Speed – This metric measures the average speed during running over a single day.
5. Number of Bouts – This metric measures the total number of discrete periods of wheel running activity over a single day.
6. Mean Bout Distance – This metric measures the mean distance of all bouts recorded on a given day.
7. Median Bout Distance – Same as above, except median.
8. Mean Bout Duration – This metric measures the mean duration of all bouts recorded on a given day.
9. Median Bout Duration – Same as above, except median.
10. Mean Interbout Duration – This metric measures the mean amount of time recorded between individual running bouts on a given day.
11. Median Interbout Duration – Same as above, except median.
12. Short-Bout Proportion – This metric measures the proportion of all bouts on a given day that were short bouts, based on a user-defined threshold previously inputted in the settings panel.
13. Active-Cycle Proportion – This metric measures the proportion of all bouts on a given day that were completed during an animal’s active cycle, a user-defined threshold.

**Figure 7.**
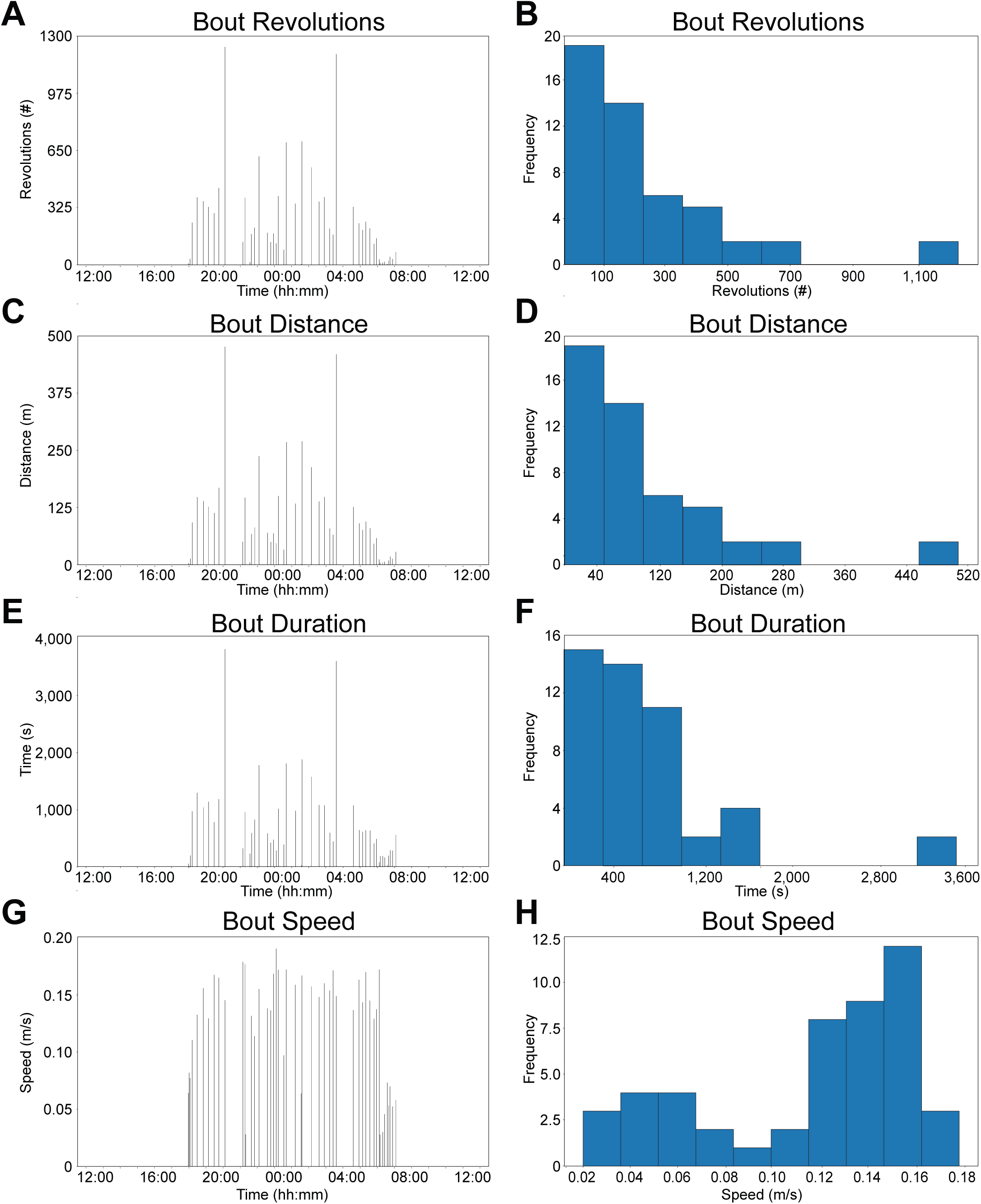
REVS day level outputs. The outputs reflect the bout timing and bout histogram outputs from the explore panel. These outputs allow the user to visualize the exact timing of each bout and its associated revolution count (**7A**), distance (**7C**), duration (**7E**), and speed (**7G**) for a specific animal on a specific day. Each line represents one bout. Users can also measure the frequency or proportion of bouts in the form of a histogram (**7B**, **7D**, **7F**, **7H**). Associated data is available for export via the export panel.

**Figure 8.**
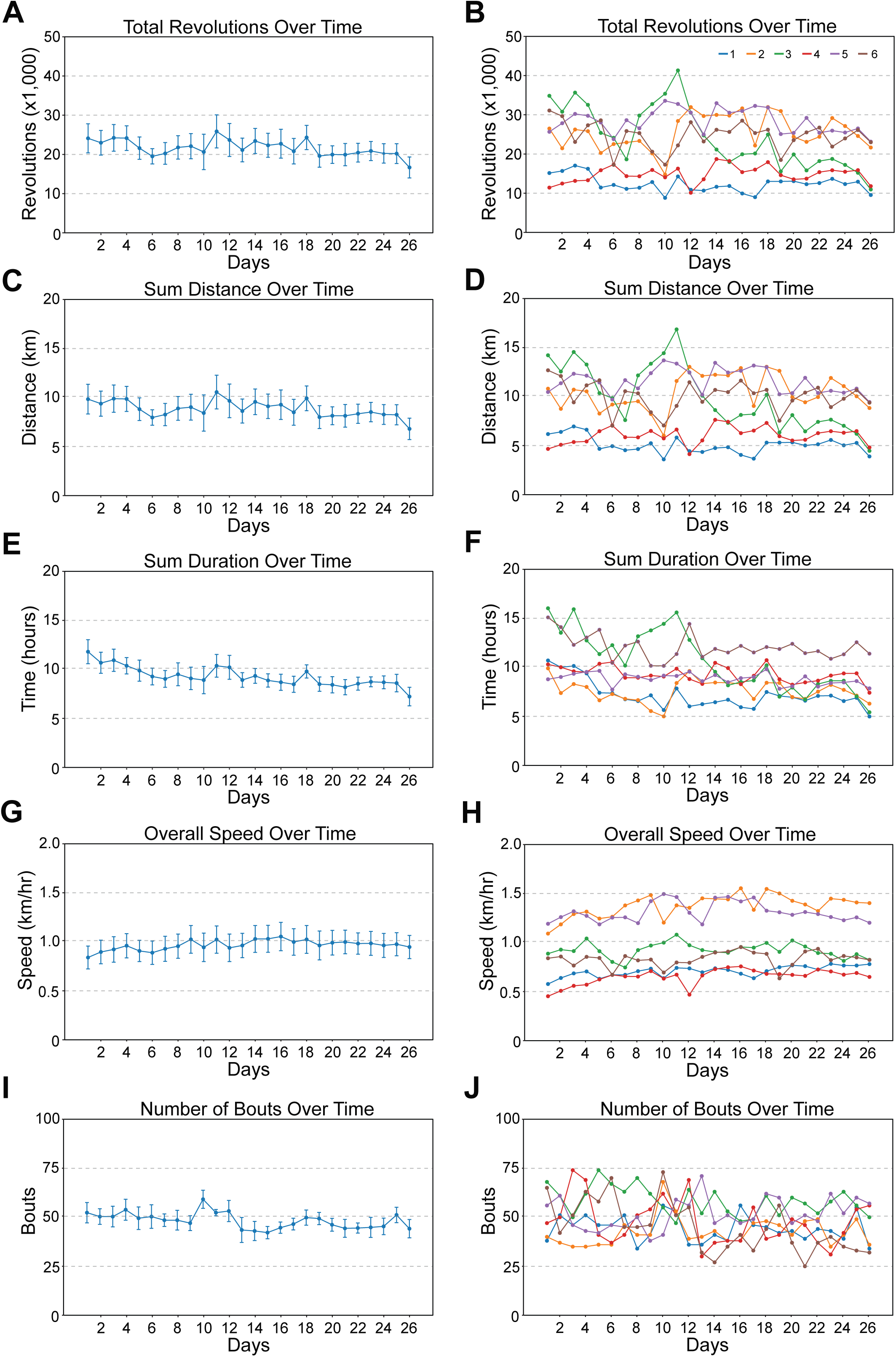
REVS day level metrics. The outputs above reflect the population means and individual animal day-level metrics from the explore panel. These outputs allow the user to visualize the mean and individual measurements respectively for revolutions **(8A, 8B)**, distance **(8C, 8D)**, duration **(8E, 8F)**, speed **(8G, 8H)**, and bout number **(8I, 8J)**. Associated raw data is available for export in the export panel.

###### 2.4.3.3.1 Bout Timing and Bout Measurement Histograms

The third and fourth graph functionality of the Explore panel allows the user to visualize bout timing and histograms of bout measurements (Fig. 9). The bout timing visualization shows the revolutions, distance, duration, or speed of every recorded bout with time stamps for one animal on a given day. The histogram option allows users to interpret the distribution of these bout metrics. As with the grouped and individual mean data, individual graphs and/or all raw data can be exported for further analyses.

**Figure 9.**
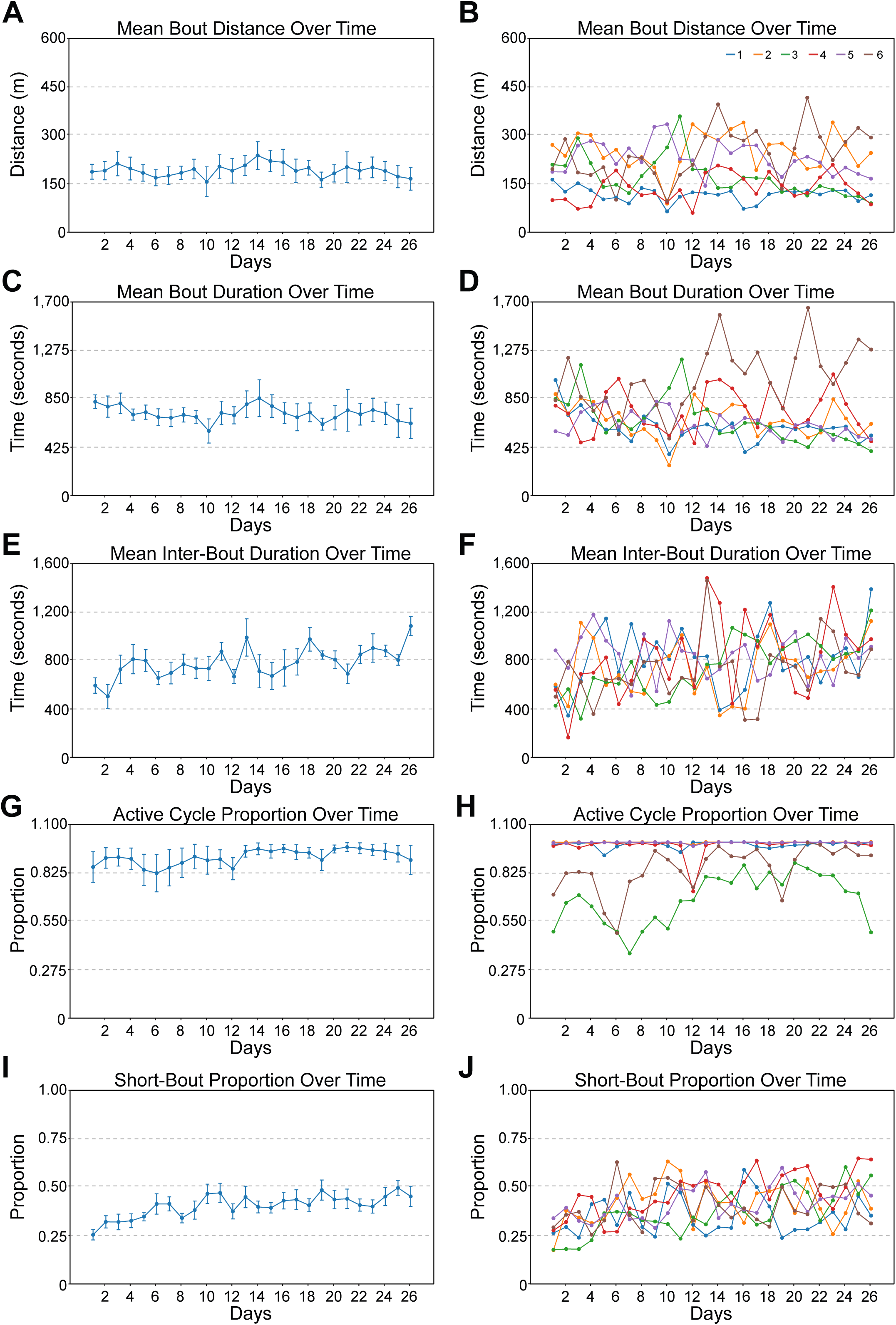
REVS bout level metrics. The outputs above reflect the population means and individual animal bout-level metrics from the explore panel. These outputs allow the user to visualize the mean and individual measurements respectively for bout distance **(9A, 9B)**, duration **(9C, 9D)**, inter-bout duration **(9E, 9F)**, active cycle proportion **(9G, 9H)**, and short-bout proportion (**9I**, **9J**). Associated raw data is available for export in the export panel.

##### 2.4.3.4 Export Panel

Finally, the export panel allows users to view and modify their metadata spreadsheet and export the entire dataset (Fig. 10). The GUI expects a JSON file which outlines the ODC data requirements, which is selected by the user in the settings panel. Based on the chosen ODC framework, the export panel will allow users to identify and add missing data for ODC compatibility. After finishing, the user can select the “Export” button to get an ODC-compatible dataset. The “Export” button also creates a data dictionary, a structured document required by ODC which lets users define and describe their variables. The data dictionary will output as a .csv, allowing the user to make final additions and adjustments.

**Figure 10.**
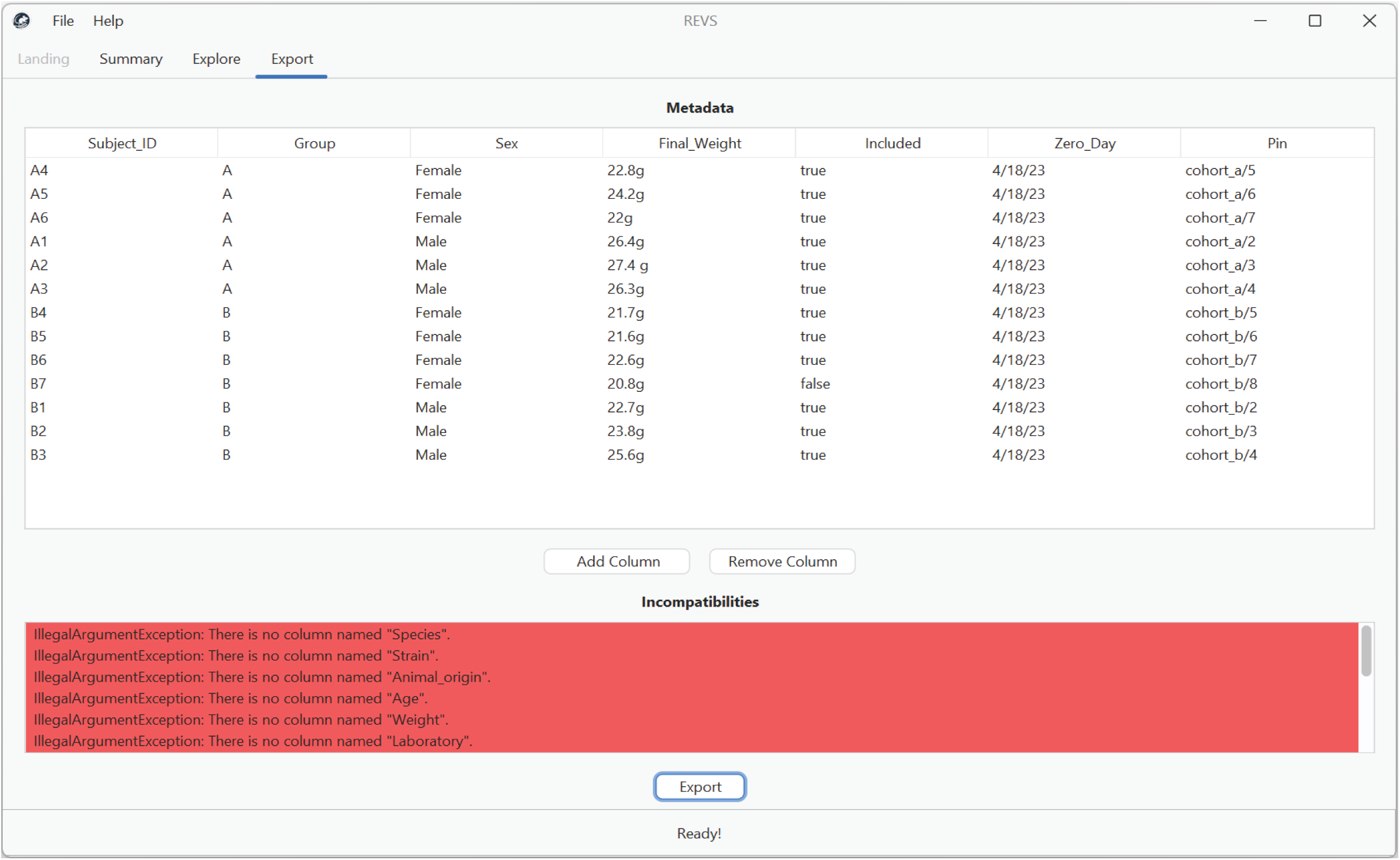
REVS export panel. The REVS export panel allows users to view their metadata spreadsheet and add columns that are necessary for ODC compatibility. When columns for ODC compatibility are missing, it will be noted in the incompatibilities dialog box. Users can still ignore these warnings and export their dataset. In addition to exporting all data, hitting the Export button also creates an ODC-compatible data dictionary.

### 2.5 Study Details

#### 2.5.1 Mice and surgery

To demonstrate the utility of the REVS pipeline, we asked whether REVS could identify lesion-induced changes and recovery signatures after partial spinal cord injury. 6 male and 7 female adult C57BL/6 mice, aged 6-8 weeks, were used in this experiment. All mice were assigned randomly to cages and given access to a runged running wheel for habituation and baseline data collection. Then all mice received a unilateral pyramidotomy at the level of the medulla as previously described (Cafferty & Strittmatter, 2006). Two days following the lesion, mice regained access to complex running wheels for four weeks. All surgical procedures and postoperative care were performed in accordance with guidelines of the Yale Animal Care and Use Committee. One female mouse was excluded from data analysis because it stopped running entirely shortly after the lesion. Running logs were collected daily and analyzed using the REVs software. A separate cohort of 6 naive C57BL/6 male and female mice were used to collect the data presented in Figs. 7-9 in the Methods section.

#### 2.5.2 REVS Settings

The following settings were manually inputted or selected in the REVS settings panel: wheel circumference, 0.408m; minimum interbout duration, 5s; short bout threshold, 120s; minimum revolution threshold, 100; day boundary, 11:30; start of active cycle, 18:00:00; start of inactive cycle, 06:00:00. ODC-SCI compatible export parameters were provided in a JSON file.

#### 2.5.2 Statistics

A within-subjects study design was used to determine the effects of pyramidotomy and rehabilitation. Specifically, we performed a repeated measures ANOVA using the baseline, day 3, and day 28 for each of the 13 REVS output metrics to identify lesion-induced differences. Analyses were restricted to animals that ran greater than 3 km at baseline, resulting in one exclusion. Two animals that became sick post-lesion fell below this threshold and were removed from analyses due to incomplete data sets. Bonferroni-adjusted *post-hoc* tests were performed between each of the days if a significant main effect was found. A p-value of 0.05 was selected as a cutoff for significance *a priori*.

## 3 Results

### 3.1 Overview of REVs hardware and software

Here, we provide a detailed overview for building an open-source, low-cost wheel-running system that is capable of tracking wheel revolution data for singly housed mice with high temporal resolution. Our system consists of an Arduino Uno R3 that uses Hall effect sensors to detect the revolutions of magnetized in-cage running wheels (Figs. 1-3). In addition to providing a system for chronic wheel revolution data collection, we also provide a user-friendly graphic user interface (GUI) which allows users to easily visualize their data and export it in formats compatible with Open Data Commons (ODC) requirements (Figs. 4-10). REVS detects wheel revolution characteristics with granular resolution. To demonstrate the utility of this new behavioral acquisition and data visualization approach we collected data from naïve adult wild type mice over a 26-day period (Figs. 7-8). REVS outputs include day level metrics that can be viewed at the group or individual animal level. Precise temporal resolution of activity at the bout level can be explored within individual animals on any specific day. These detailed metrics allow for the detection of disease or injury-induced alterations in locomotor characteristics.

### 3.2 Effect of Unilateral Pyramidotomy and Rehabilitation on individual wheel running metrics

To explore the impact of partial spinal cord injury on locomotor function, we performed unilateral pyramidotomies (uPyX) on adult mice and compared their wheel running metrics over 28 days compared to a baseline, uninjured timepoint (Fig. 11). We detected a change from baseline in 10 of the 13 metrics using repeated measures ANOVAs, summarized in Table 1. For whole-day metrics, we detected a deficit between baseline and day 3 for total revolutions (F=21.4, p<0.001), total distance (F=21.4, p<0.001), total duration (F=4.3, p=0.008), overall speed (F=52.9, p<0.001), and short-bout proportion (F=10.4, p=0.003). Additionally, deficits were detected in the following bout-level metrics: mean bout distance (F=20.7, p<0.001), median bout distance (F=13.9, p<0.001), mean bout duration (F=9.0, p<0.001), median bout duration (F=12.8, p<0.001), and median interbout duration (F=9.4, p=0.007). There were no deficits for number of bouts, active cycle proportion, and mean interbout duration.

**Figure 11.**
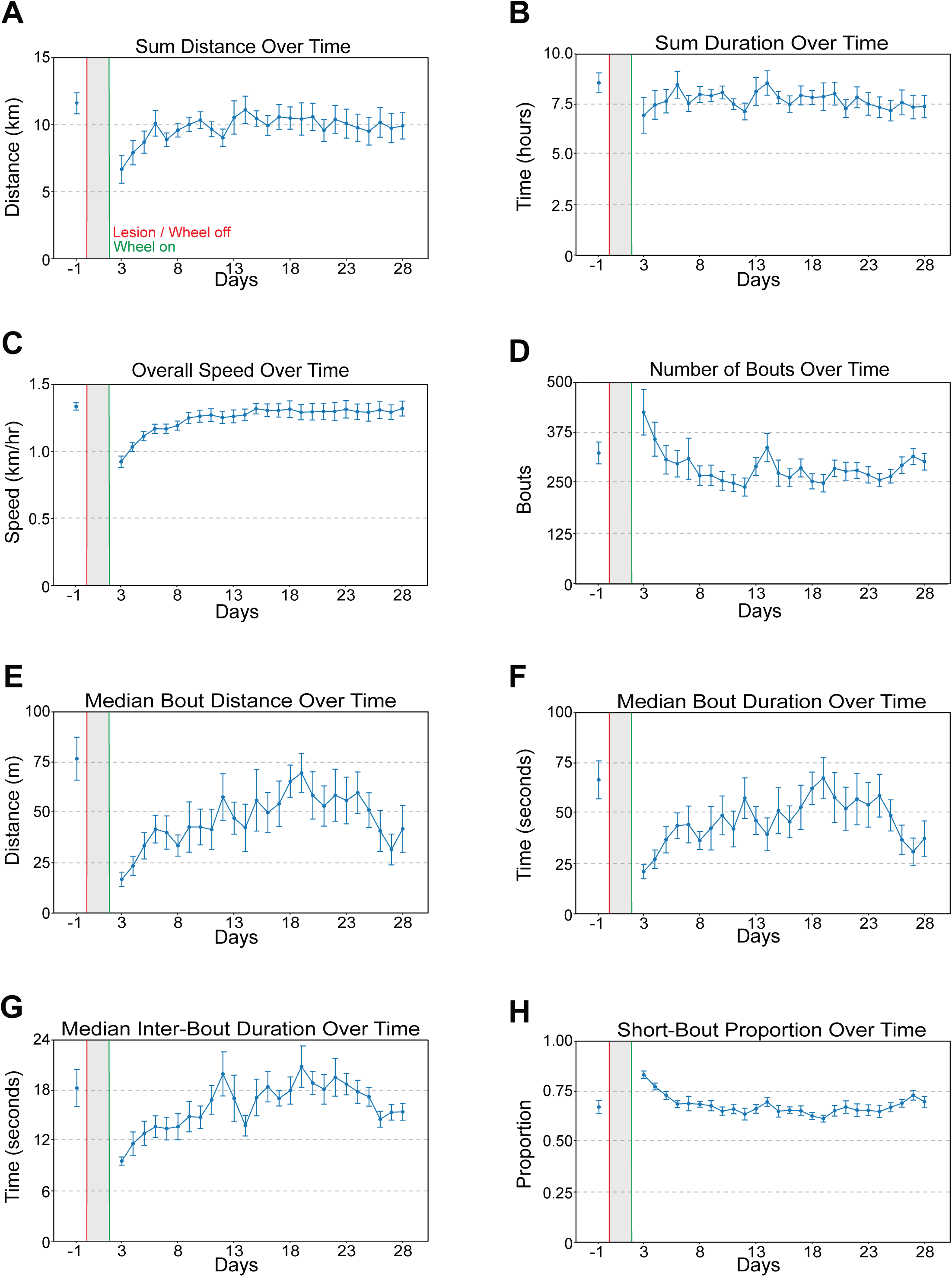
Effects of unilateral corticospinal tract lesion on REVS metrics. Unilaterally lesioned mice were given complex wheels and their wheel-running behavior was measured for four weeks. Red lines indicate the day of lesion (day 0). Wheels were removed for two days and replaced at day 3 (green lines). Mice then experienced 25 consecutive days of running. Affected metrics include total distance **(11A)**, duration **(11B)**, speed **(11C)**, bout number **(11D)**, median bout distance **(11E)**, median bout duration **(11F)**, median interbout duration **(11G)**, and short-bout proportion **(11H)**.

**Figure 12.**
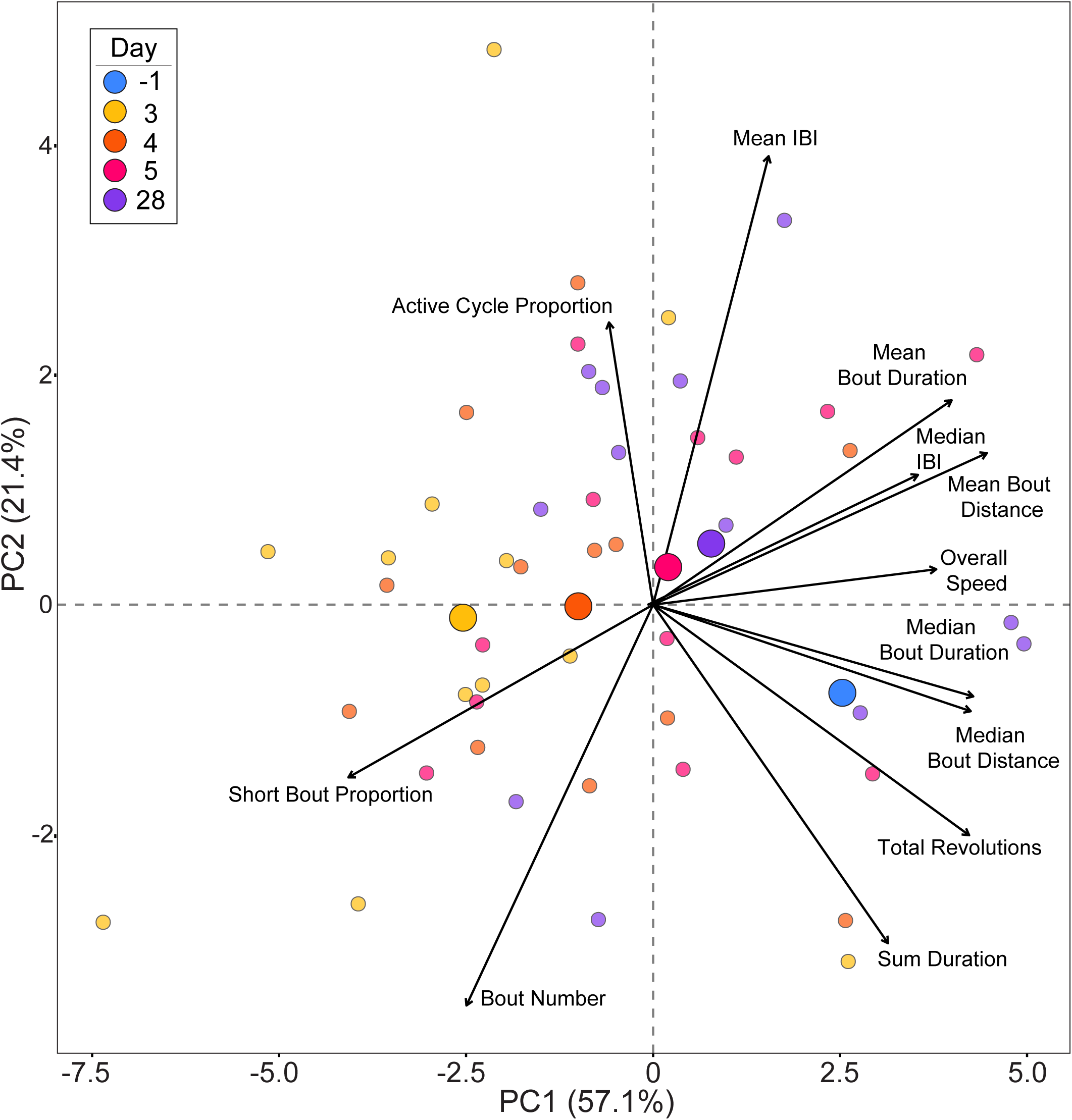
Principal component analysis reveals lesion recovery profile. The thirteen REVS metrics for each animal were normalized to their baseline performance and used in principal component analysis (PCA). The PCA revealed an acute moderate recovery along the PC1 axis, which was most strongly driven by bout characteristics including duration and speed. There was no recovery along the PC2 axis, which was most strongly driven by bout timing and number.

**Table 1.**
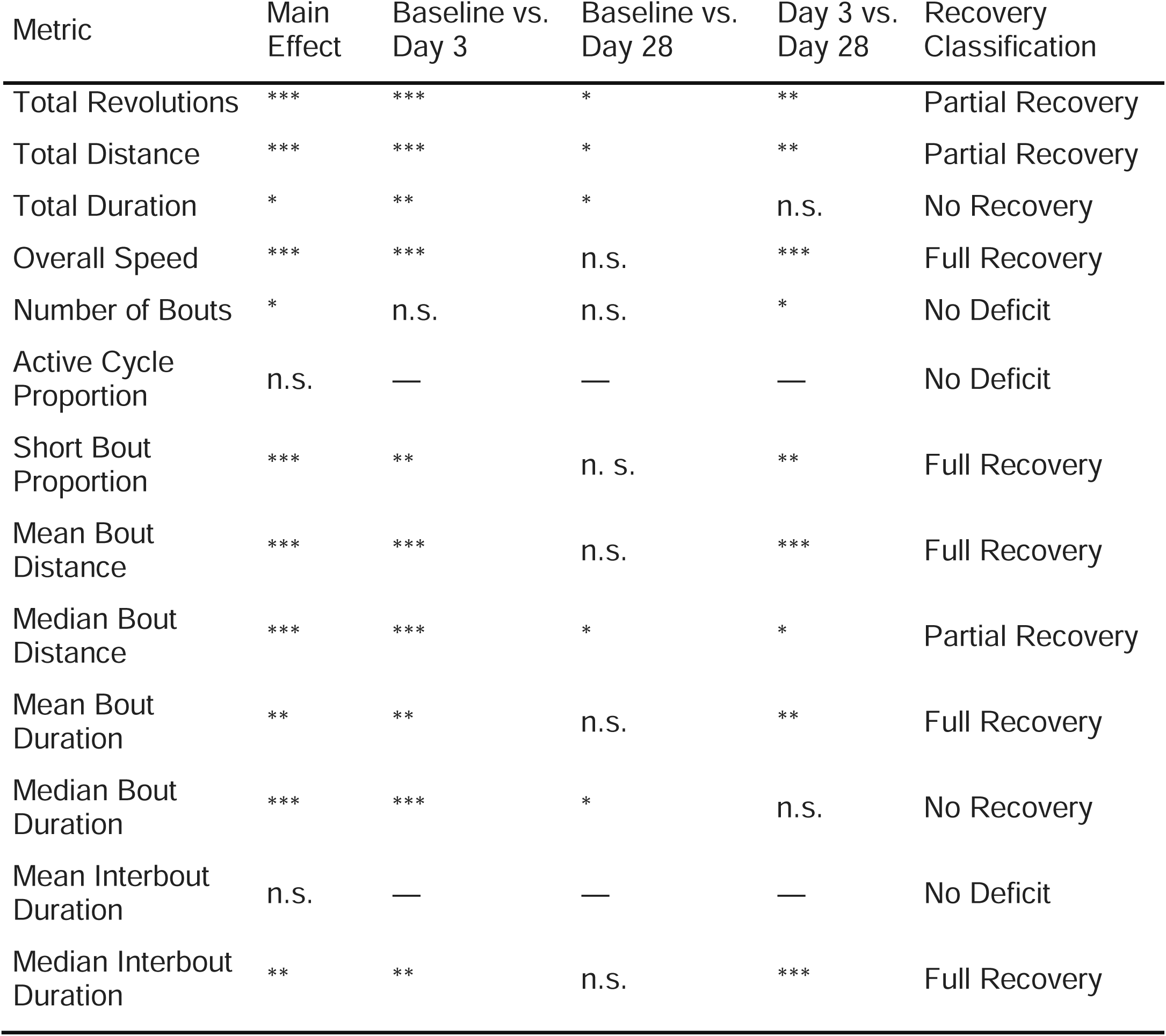
Functional metric changes over time after unilateral corticospinal tract lesion. The main effect of lesion and rehabilitation is reported for each of the thirteen REVS metrics (repeated measures ANOVA). Eleven metrics showed a significant main effect. Bonferroni corrected *post hoc* testing revealed that ten metrics showed a significant deficit between baseline and day three, while two metrics showed no deficit. The ten deficit metrics were classified further into full recovery, partial recovery, or no recovery based on the pattern of their significant differences between baseline versus day twenty-eight, and day three versus day twenty-eight. *=p<0.05, **=p<0.01, ***=p<0.001, n.s. = not significant.

| Metric | Main Effect | Baseline vs. Day 3 | Baseline vs. Day 28 | Day 3 vs. Day 28 | Recovery Classification |
| --- | --- | --- | --- | --- | --- |
| Total Revolutions | *** | *** | * | ** | Partial Recovery |
| Total Distance | *** | *** | * | ** | Partial Recovery |
| Total Duration | * | ** | * | n.s. | No Recovery |
| Overall Speed | *** | *** | n.s. | *** | Full Recovery |
| Number of Bouts | * | n.s. | n.s. | * | No Deficit |
| Active Cycle Proportion | n.s. | — | — | — | No Deficit |
| Short Bout Proportion | *** | ** | n. s. | ** | Full Recovery |
| Mean Bout Distance | *** | *** | n.s. | *** | Full Recovery |
| Median Bout Distance | *** | *** | * | * | Partial Recovery |
| Mean Bout Duration | ** | ** | n.s. | ** | Full Recovery |
| Median Bout Duration | *** | *** | * | n.s. | No Recovery |
| Mean Interbout Duration | n.s. | — | — | — | No Deficit |
| Median Interbout Duration | ** | ** | n.s. | *** | Full Recovery |

Of the 10 metrics for which deficits were detected, we classified them into recovery profiles of no recovery, partial recovery, and full recovery depending on their pattern of significance differences between day 3 and day 28, and baseline and day 28. Metrics that had no difference between day 3 and day 28 were classified as having no recovery; metrics that had a difference between day 3 and day 28 were classified as partial or full recovery depending on whether the day 28 timepoint was different from baseline.

Using this classification, two metrics had no recovery: total duration (D3 v. D28, p=0.55) and median bout duration (D3 v. D28, p=0.58). Three metrics had a partial recovery: total revolution and total distance (D3 v. D28, p=0.008), and median bout distance (D3 v. D28, p=0.03). Five metrics had a full recovery: overall speed (baseline v. D28, p=0.34), short-bout proportion (baseline v. D28, p=0.42), mean bout distance (baseline v. D28, p=0.13), mean bout duration (baseline v. D28, p=0.23), and median interbout duration (baseline v. D28, p=0.30). These data demonstrate the capacity of REVS to identify lesion and rehabilitation-induced changes in locomotor characteristics over time.

### 3.3 Principal components analysis of uPyX Recovery Profile

To further explore the recovery profile of uPyX mice, we performed a principal components analysis of the 13 wheel running metrics including days 3-5, for which the recovery occurred, and day 28, the final time point in the study. The strongest predictors of the first principal component (PC1, 57.1%) were mean bout distance (eigenvector = 0.36), median bout duration (0.34), median bout distance (0.34), total revolutions and distance (p=0.34), and short-bout proportion (-0.32). Together, these loadings represent the intensity of activity during bouts.

In contrast, the strongest predictors of the second principal component (PC2, 21.4%) were mean interbout duration (0.51), number of bouts (-0.45), total duration (-0.38), and active cycle proportion (0.32). Together, these loadings represent the frequency and timing of bouts.

Overall, both axes demonstrate a shift away from the baseline timepoint immediately after injury, but especially PC1. Over the course of days 3-5, mice sharply move back toward the baseline time point along the PC1 axis, which quickly plateaus. By day 28, there is still a considerable distance from baseline, consistent with the partial recovery observed in individual metrics. By contrast, PC2 fails to trend back toward the baseline timepoint throughout the recovery process. Together, this reveals that although mice undergo alterations in both the intensity and frequency of activity post-injury, it is primarily changes in intensity that are a hallmark of rehabilitation-associated recovery. The ability to explore the recovery profile through PCA is afforded by the level of detail REVS provides on the locomotor characteristics.

## 4. Discussion

### 4.1 REVS Overview, Strengths, and Limitations

The use of running wheels in animal studies is longstanding and widespread across a variety of fields of study, such as exercise physiology, circadian rhythm, cognition and addiction, and nervous system injury and rehabilitation (Battistuzzo et al., 2012; Guo et al., 2019). However, the amount of data acquired from studies involving wheels can vary substantially – from no data regarding wheel running behavior at all, to simple measurements of distance and speed, to more advanced quantifications like activity diagrams. Applications range in use but mostly fall into one of two categories: in-house customized set-ups and for-purchase hardware and software, each with its own benefits and drawbacks. While in-house approaches are a more affordable, accessible, and customizable option, they may only provide simple details about wheel running characteristics like total distance ran which require the development of data processing pipelines. In contrast, while for-purchase hardware and software often provide more detailed and streamlined wheel behavior outputs, they are often cost-prohibitive and their components non-modifiable. We sought to provide an alternative with the benefits of both: low-cost, modifiable hardware combined with a detailed, open-source, and user-friendly data analysis, visualization and export program.

REVS has several advantages compared to previously developed technology. First, REVS takes an experimental question-focused approach by allowing the experimenter to select the type of wheel used. Because REVS uses Hall effect sensors attached to individual wheels to detect revolutions, any wheel can be used if it can fit in vivarium-approved cages. In spinal cord injury research, injuries can often make it difficult for rodents to use wheels with rungs (Engesser-Cesar et al., 2005). This renders the use of many for-purchase hardware unusable because their wheels are typically standardized and non-modifiable. In such cases, REVS allows for the use of solid-bottomed, smooth wheels. In the present study, we used a partial spinal cord injury model that particularly affects fine motor control (Starkey et al., 2005). Accordingly, to target mice for rehabilitation we modified our wheels to have rough, sanded rungs with a complex pattern. Notably, our system also makes it possible to switch out wheel types throughout the experiment, allowing for more complex research questions and the use of regular or complex upright wheels. This stands in contrast to previously developed open-source wheel-tracking approaches, which are designed for the use of a disk-running wheel (Deitzler et al., 2022; Edwards et al., 2021; Godfrey et al., 2022; Terstege & Epp, 2024; Zhu et al., 2021).

A second advantage of REVS over other hardware and software is the level of detail it provides for the user. In addition to measuring standard wheel running metrics like revolutions, distance, and speed, REVS measures 9 additional metrics that can be useful in assessing impacts of injuries and interventions. These include the number of bouts, mean and median distances and duration of bouts, the mean and median interbout durations, and the proportions of bouts that were short/long or during the active/inactive cycle. All 13 of these metrics can be easily viewed in REVS software at the group and individual animal level. Moreover, group-level visualization is easily modifiable to stratify or collapse across multiple grouping keys, such as sex and treatment. Visualizations can be simply exported with associated data. Finally, REVS allows users to explore the individual bout-level data for a specific subject ID on a specific day with two distinct approaches: millisecond-level bout timing graphs which provide the number of revolutions, distance, speed, or duration for each bout, or a histogram which provides distributions of the revolutions, distances, speeds, or durations of bouts. Full data for each of these approaches can be exported to ask questions at the group and/or individual level. These advancements of REVS provide advantages compared to previously developed open-source options, which have limited metrics, lack a GUI entirely, and and/or require the development of a data analysis pipeline (Deitzler et al., 2022; Edwards et al., 2021; Godfrey et al., 2022; Terstege & Epp, 2024; Zhu et al., 2021).

A final advantage of the REVS software is the built-in capacity to easily export all data in a format consistent with the most prominent open data framework, the Open Data Commons (ODC) (Chou et al., 2022; Heath et al., 2021; Torres-Espín et al., 2022). Transparency of published data is often lacking, despite a growing recognition of the importance of open-source data and the requirement by some international-funding bodies to provide it. REVS allows the user to upload a list of ODC requirements, which the program will require as input in the Export panel. These metadata can easily be added in an editable table within the panel. Upon export of data, REVS will simply merge the collected data with required ODC metadata. Crucially, this approach will significantly decrease the workload required to make data ODC-compatible, which can be cumbersome when data is collected with missing ODC requirements. In addition to exporting data in an ODC-compatible format as a .csv, the export panel will also output a partially completed data dictionary, or glossary of relevant experimental variables. To the authors’ knowledge, REVS is the first out of all previously available wheel revolution hardware and software to provide a fully integrated ODC-compatible data export.

REVS does have some limitations compared to other options. First, REVS is most compatible when animals are singly housed – raising the possibility of social isolation as a behavioral confound (Mayr et al., 2020). In our experience, this did not prohibit mice from significant wheel usage. In our setup, singly housed mice were placed in cages directly adjacent to at least one other cage, facilitating communication between mice. Of note, it is not a requirement that animals be singly housed: rodents can be group housed, and cage-level data can be collected. However, previous literature suggests that group-housed mice may compete for wheel usage and experience increased aggression (Akre et al., 2011; Swetter et al., 2011). To avoid singly housing mice, RFID technology could be used to allot the proportion of certain wheel running metrics, such as total distance, to group housed animals (Habedank et al., 2022). Finally, REVS does not have the ability to lock wheels remotely at any time, something that some software packages include (Bivona & Poynter, 2021; Mayr et al., 2020). It does, however, remain possible to manually lock wheels at certain time points if an experimenter chooses.

### 4.2 Interpretation of Results

To demonstrate its utility, we performed an experiment to test whether REVS could identify lesion-induced deficits and hallmarks of recovery in a mouse model of partial spinal cord injury. Specifically, we acquired a baseline time point for male and female, adult C57BL/6 mice and then performed unilateral corticospinal tract injury (pyramidotomy). These lesions result in significant and lasting compromise of fine motor control (Starkey et al., 2005). We expected to find an impairment in at least some of REVS’ 13 whole-day metrics, given that we used complex wheels that require fine motor coordination. Indeed, we observed at least a transient deficit in 10 of the 13 metrics (Fig. 11). Interestingly, the 10 metrics that showed deficits did not all exhibit the same recovery profile. For example, only 5 metrics – overall speed, mean and median bout distance, mean bout duration, and median interbout duration – made a full recovery by day 28 from their day 3 impairment. 3 metrics recovered partially including median bout distance, total distance, and total revolutions; while 2 metrics, total duration and median bout duration, failed to recover by the end of the testing period. Additionally, 3 metrics showed no impairment from the lesion, including total number of bouts, active cycle proportion, and mean interbout duration. A Principal Component Analysis of our 13 metrics revealed that PC1 loadings, which were representative of bout quality and intensity characteristics were most associated with a recovery while PC2 loadings representative of bout timing were not.

Together, these analyses reveal that REVS facilitates the detection of subtle changes in motor function in the context of injury and rehabilitation. These results underscore the importance of identifying and measuring subtle changes in behavior in mouse models of injury and disease. We speculate that REVS may be able to identify unique constellations of behavioral recovery associated with different levels and types of injury, or levels of treatment and interventions in ways that other behavioral measures may fail to observe or detect. Therefore, REVS is not only a tool for administering exercise and rehabilitation in studies of mouse models of physiology and injury but is also a tool for the detection of unique behavioral phenotypes over time. REVS as a tool for behavioral phenotype detection will provide a useful complement to studies of rehabilitation that often use other behavioral tasks to quantify recovery and to provide further insight into task-specific and non-task specific recovery profiles.

### 4.3 Summary

In summary, here we introduce REVS, an open-source and low-cost hardware and software for measuring rodent wheel running with highly detailed behavioral outputs. In addition to granular behavioral outputs, REVS also affords flexibility with regard to experimental design and streamlines the process of data analysis, visualization, export, and sharing. To demonstrate the value of REVS, we used it to identify a unique profile of behavioral impairments and recovery in a mouse model of partial spinal cord injury. However, we are excited about the broad applicability of REVS, which can be used to detect detailed data about mouse locomotor behavior not only in other studies of injury and rehabilitation, but also beyond.

## Declaration of generative AI and AI-assisted technologies in the writing process

During the preparation of this work the author(s) used ChatGPT based on GPT-4o to improve the clarity and readability of the manuscript. After using this tool/service, the author(s) manually reviewed and edited proposed content and clarity changes as needed and take(s) full responsibility for the content of the publication.

## Glossary

Voluntary Wheel Running: A naturalistic, self-initiated locomotor activity used to assess behavior or promote recovery.
Spinal Cord Injury (SCI): Damage to the spinal cord that impairs motor and/or sensory function, often modeled in rodents.
Pyramidotomy: A lesion model in which the corticospinal tract is severed at the level of the medullary pyramids, resulting in deficits in fine motor control
Behavioral Phenotyping: Characterization of specific behavioral patterns, here in the context of nervous system injury
Bout: A discrete period of wheel running activity separated by periods of inactivity
Hall effect sensor: a device that detects magnetic fields and is used in REVS to measure wheel revolutions
Interbout Interval: The time between two consecutive running bouts.
Active Cycle: Also known as the night cycle, or the period of the day in which mice are awake and active
Open Data Commons (ODC): A framework promoting standardized metadata and formats for scientific data sharing
Data Dictionary: A structured reference that defines variables and metadata used in a dataset.
Metadata: Contextual information (e.g., animal ID, treatment, timepoints) necessary for interpreting a dataset.
Principal Component Analysis (PCA): A statistical method used to reduce data dimensionality and identify patterns in multivariate datasets

## Author contributions: CRediT

**James Bonanno:** Conceptualization, data curation, formal analysis, methodology, investigation, project administration, visualization, validation, writing – original draft; **Ciara O’Brien:** Data curation, methodology, software, validation; **William. B.J. Cafferty:** Conceptualization, methodology, resources, supervision, validation, writing – review and editing.

## Data statement

All data from the methods and results sections are publicly available on the ODC-SCI.

### Acknowledgements

The authors would like to acknowledge Peter O’Brien for his expert technical assistance in prototyping the REVS hardware.

